# Time-series expression profiling of sugarcane leaves infected with *Puccinia kuehnii* reveals an ineffective defense system leading to susceptibility

**DOI:** 10.1101/584276

**Authors:** Fernando Henrique Correr, Guilherme Kenichi Hosaka, Sergio Gregorio Perez Gomez, Mariana Cicarelli Cia, Claudia Barros Monteiro Vitorello, Luis Eduardo Aranha Camargo, Nelson Sidnei Massola Júnior, Monalisa Sampaio Carneiro, Gabriel Rodrigues Alves Margarido

## Abstract

*Puccinia kuehnii* is an obligate biotrophic fungus that infects sugarcane leaves causing a disease called orange rust. It spread out to other countries resulting in reduction of crop yield since its first outbreak in Australia. One of the knowledge gaps of that pathosystem is to understand the molecular mechanisms altered in susceptible plants by the stress induced by *P. kuehnii*. Here we investigated changes in temporal expression of transcripts in pathways associated with the immune system. To achieve this purpose, we used RNA-Seq to analyze infected leaf samples collected at 0, 12, 24, 48 hours after inoculation (hai), 5 and 12 days after inoculation (dai). A *de novo* transcriptome was assembled with preprocessed reads, filtering out potential fungal sequences to focus on plant transcripts. Differential expression analyses of adjacent time points revealed substantial changes at 12, 48 hai and 12 dai, coinciding with the events of spore germination, haustoria post-penetration and post-sporulation. During the first 24 hours, a lack of transcripts involved with resistance mechanisms was revealed by underrepresentation of hypersensitive and defense responses. However, two days after inoculation, upregulation of genes involved with immune response regulation provided evidence of some potential defense response. Events related to biotic stress and phenylpropanoid biosynthesis pathways were predominantly downregulated in the initial time points, but expression was later restored to basal levels. Similar waves of expression were apparent for sets of genes in photosynthesis and oxidative processes, whose initial repression could avoid production of signaling molecules. Their subsequent upregulation possibly restored carbohydrate metabolism, ensuring pathogen nutritional requirements were met. Our results support the hypothesis that *P. kuehnii* initially suppressed sugarcane genes involved in plant defense systems. Late overexpression of specific regulatory pathways also suggests the possibility of an inefficient recognition system by a susceptible sugarcane genotype.

## Introduction

Sugarcane is currently cultivated in 27 million hectares worldwide, 52% of which are located in America, more expressively in South America (11 million ha). Brazil is the main producer, responsible for 768.68 million tonnes (FAOSTAT, 2016). The term sugarcane includes at least six species of the genus *Saccharum* [1, 2]. The ancestor species are *Saccharum robustum* and *S. spontaneum*, which present variable ploidy levels with chromosome number ranges of 2n = 60 - 170 and 2n = 36 - 128, respectively [3, 4]. The octoploid (2n = 80) *S. officinarum* is an ancient cultivated sugarcane with high sugar content that formed the group of noble cultivars [3, 5, 6]. Sugarcane plants currently used for industrial purposes are modern cultivars derived from interspecific crosses, mainly between *S. officinarum* and *S. spontaneum*. As a consequence of successive backcrosses, the majority of chromosomes in homology groups are from *S. officinarum*, with a less frequent occurrence of *S. spontaneum* chromosomes, some recombinant chromosomes and frequent aneuploidy [7, 8].

Sugarcane is important for sugar and ethanol production and, moreover, new perspectives emerged because of its capacity for energy production and due to new uses for the industrial residues [9–12]. Despite its economic importance, sugarcane production is harmed by insect and nematode attacks, weed competition and diseases, leading to losses in productivity [13]. Among the diseases currently affecting sugarcane crops, three important biotrophic fungi stand out: i) *Puccinia melanocephala*, responsible for brown rust; ii) *Puccinia kuehnii*, causal agent of orange rust; and iii) *Sporisorium scitamineum*, which causes smut. Both rusts affect leaf tissue, while smut affects meristem development. Particularly, more attention was given to orange rust following an outbreak in Australia roughly eighteen years ago [14]. Occurrences were then reported in America in the last ten years [15, 16], but noticeably with smaller diversity of isolates than that observed in the Eastern hemisphere, suggesting a spread of particular strains from Asia and Australia [17]. The impacts of *P. kuehnii* on sugarcane yield have been associated with variation in photosynthesis, because there are considerable changes in leaf physiological measures in susceptible cultivars where symptoms are evident [18, 19].

Biotrophic pathogens require living host tissues to capture nutrients for their development. The establishment of biotrophic fungi consists in spore adhesion, germination on plant surface and formation of appressoria, which are the penetration structures. Later, the spread of fungal cells occurs by different modes of colonization. Obligate biotrophs, like the species of the genus *Puccinia*, initially grow hyphae inside the host tissues, moving to intracellular colonization by haustoria, side branches from these hyphae [20, 21]. Haustoria act as an interface to acquire nutrients from the host and to deliver effectors [20, 22]. Effectors are essential molecules secreted by pathogens to manipulate host metabolism to control the plant immune system. They may be recognized by plant resistance proteins, which is generally followed by an oxidative burst and promotion of the hypersensitive response. The relevance of the latter is that it induces cell death to restrict pathogen development, ensuring the non-capture of nutrients and water. Resistance gene products also activate hormone-dependent signaling, by salicylic (SA) or jasmonic acid (JA), which regulates expression of other defense response genes [21, 23].

Plant infection is a dynamic process and, at the RNA level, the expressed genes of host and pathogen reflect the so-called interaction transcriptome [24]. Through this concept, it is possible to capture the profiles of transcripts with relevant roles, such as the members of the signaling pathways which prevent disease establishment. Techniques that measure changes in RNA levels such as RNA Sequencing (RNA-Seq) are used to detect differentially expressed genes. This methodology is based on the development of Next-Generation Sequencing (NGS) technologies, enabling the study of variations in RNA composition between samples from distinct experimental conditions. RNA-Seq was used in sugarcane to provide molecular information about transcripts from a range of superior genotypes [25], understanding changes in lignin content [26] and the development of organs [27]. With regard to pathogenic interactions, RNA-Seq has the sensitivity to capture both pathogen and host transcripts, not only providing transcripts catalogs, but also changes in gene expression levels due to the infection process [28]. In plants, transcriptome profiling studies are covering pathosystems in annual [29–33] and perennial species [34–38]. Recently, studies concerning the interaction of sugarcane with the fungus *S. scitamineum* [39–41], with the bacteria *Acidovorax avenae* subsp. *avenae* [42] and with the bacteria *Leifsonia xyli* subsp. *xyli* were published [43].

Knowledge about the molecular mechanics of infection in sugarcane is important and there are efforts in identifying molecular markers for disease resistance [44–46]. Efforts were taken for developing molecular markers associated with resistance against both sugarcane rusts. For brown rust, markers associated with the *Bru1* QTL have been used in sugarcane research [44, 47, 48], mainly for breeding purposes [44], as well as for investigations about its occurrence in genotype panels [47]. With regard to orange rust, cultivars have shown distinct responses when infected with *P. kuehnii* strains [49].

Primarily, studies in orange rust focused on molecular detection of *P. kuehnii* by using primers that amplify ITS regions [17], the characterization of pathogen developmental structures during infection [50] or the assessment of infection degree in sugarcane cultivars [18, 19]. Yang and collaborators [46] detected three QTL associated with resistance against orange rust. They also identified putative resistance genes based on sugarcane transcripts with orthologs in the sorghum genome. To that end, they searched for homology with leucine-rich-repeat (LRR) and nucleotide binding (NB) sites using five combined databases. One identified gene allowed the development of a marker significantly associated with orange rust resistance. Also, non-inoculated susceptible and resistant genotypes were grouped based on metabolite fingerprinting [51]. However, there are no previous studies gauging gene expression changes as a result of rust infections in sugarcane. To our knowledge, there is no information about high-throughput molecular analyses during orange rust establishment in sugarcane. Our work thus aims to study a time series of transcriptional changes in susceptible sugarcane infected by *P. kuehnii*.

## Materials and methods

### Plant Material

Plants of the sugarcane cultivar SP89-1115, susceptible to orange rust [19, 49, 52], were grown in 500 mL plastic cups containing sterilized substrate in a greenhouse during 40 days and fertilized with an ammonium sulfate solution (30 g/L) ten days before inoculation. One day before inoculation, spores of *P. kuehnii* were obtained using a vacuum pump (Prismatec 101) from the abaxial side of infected leaves from Centro de Tecnologia Canavieira (Piracicaba, São Paulo, Brazil) that contained open pustules. A solution was prepared by homogenizing the spores for 30 minutes in distilled water, with a final concentration of 10^3^ spores/mL. Approximately 4 mL of this solution was sprayed over the plants, preferably reaching the abaxial side of leaves, and they were kept at 22 °C for 24 h in a dew chamber. After this period, they were moved to a growth chamber at 25 °C with a photoperiod of 12 h light/12 h dark. A more detailed description is available in the work developed by Gomez [50]. For molecular analyses, five biological replicates of +1 leaves were taken at 0 h, 12 hours after inoculation (hai), 24 hai, 48 hai, 5 days after inoculation (dai) and 12 dai.

### RNA extraction and sequencing

Total RNA from three biological replicates of each time point was extracted with the RNeasy Plant Mini Kit (Qiagen) in a final volume of 60 *µ*L. We assessed the quality and quantity of the RNA via electrophoresis on 2% agarose gel, ND-1000 spectrophotometer (NanoDrop) and 2100 Bioanalyzer (Agilent Technologies). Next we used aliquots to prepare mRNA-Seq indexed-libraries with the TruSeq Stranded kit (Illumina). Libraries were sequenced in two lanes of an Illumina HiSeq 2500 to obtain 125 bp paired-end reads.

Quality control of this first sequencing run showed that two libraries from the initial time point did not yield large numbers of high quality mRNA reads. Hence, we extracted RNA from two different biological replicates of the same time point and followed the same procedures to obtain sequencing libraries. In a second sequencing run (100 bp paired-end reads), we included these two samples and four samples that had already been successfully sequenced from 12, 48 hai, 5 and 12 dai. This strategy allowed us to measure and remove possible batch effects.

### Data processing, *de novo* assembly and functional annotation

Following a strategy similar to Hoang and colleagues [53], adapter sequences and low quality bases were removed with Trimmomatic [54], using windows with a minimum average Phred quality score of 20. The first 12 bases of each read were removed and only reads with at least 75 bases were kept. We used sortMeRNA [55] to filter contaminant rRNA, by aligning the reads to prokaryotic and eukaryotic rRNA databases, then selecting the remaining reads for posterior analyses.

Transcriptome *de novo* assembly was performed using Trinity 2.5.1 [56] using the default k-mer size of 25. We set the maximum number of reads to combine into a single path (*–max_reads_per_graph*) to 3,000,000. Minimum percent identity (*–min_per_id_same_path*) and maximum differences between two paths (*–max_diffs_same_path*) to merge into one were set to 90 and 10, respectively. Finally, minimum contig length (*–min_contig_length*) used for the assembly was set to 300. We established these values based on our previous analyses to minimize transcript fragmentation. We also evaluated the integrity of assembled transcripts by comparing with *Sorghum bicolor* coding sequences, the distribution of contig lengths and N50 statistics. The final transcriptome comprised the longest isoform of each transcript.

To remove potential transcripts from *P. kuehnii*, we compared the assembled transcripts with *Viridiplantae* and *Fungi* protein sequences in the Swiss-Prot database using BLASTX [57]. Transcripts matching *Fungi* sequences but not plants were discarded. Next, the remaining transcriptome was compared with the non-redundant (nr) protein database using BLASTX with an e-value cutoff of 10^−6^, considering up to 20 homologous hits. The annotation was complemented with domain and protein family assignments using InterProScan and all InterPro Consortium databases. We also used BLAST2GO [58] to map and assign Gene Ontology (GO) terms to transcripts. An additional functional annotation was performed with Trinotate [59], comparing the assembled sequences with UniProt proteins and Pfam protein domains [60].

### Differential expression and enrichment analysis

We aligned the high quality reads of each sample to the *de novo* assembled transcriptome using HISAT2 [61], with the option *–no-spliced-alignment* to disable spliced alignments. Then, a custom Python script was used to create the General Transfer Format (gtf) file for the transcriptome. We quantified gene expression levels by assigning reads to transcripts and then counting fragments with featureCounts [62], independently for each sample. Next, differential expression analysis was conducted with the edgeR package v.3.20.9 [63]. We began by selecting genes with count-per-million (CPM) greater than or equal to one in at least three samples, to filter out the lowly expressed genes. Using the full factorial model with block and time effects, we observed no significant effect of the sequencing runs. Therefore, we fitted a model considering only the time effect for all subsequent analyses. Quasi-likelihood F tests were employed to detect differentially expressed genes (DEGs) [64]. We conducted five different tests, each between a pair of adjacent time points. We also compared the initial (baseline) time point and the first day after inoculation (24 hai) to verify possible influences of circadian rhythm, checking whether the results were similar to those using the 12 hai. In order to minimize false positives, we used the Benjamini-Hochberg method at a specified False Discovery Rate (FDR) of 5% for each time point comparison.

We utilized the GOseq package [65] to identify enriched GO terms, using as the background set the genes kept after filtering for low expression. We carried out functional enrichment analyses separately for each test performed in the previous step. For all adjacent comparisons the criterion to consider a term as enriched was an FDR-adjusted *p*-value less than 0.05. This method enabled us to find GO terms enriched either as overrepresented or as underrepresented, *i.e.*, terms for which the number of DEGs was greater or less than that expected by chance alone.

### Graphical representation of temporal expression of annotated DEGs in groups of processes or specific pathways

Our functional annotation included information about the Enzyme Code (EC) for some transcripts. For each pairwise comparison, we built a file containing all up and downregulated transcripts that had an associated EC number. Next, we used KEGG Mapper (https://www.genome.jp/kegg/mapper.html) to search for global metabolic maps containing DEGs.

To visualize processes and pathways as informative diagrams we used MapMan [66]. First, we obtained the appropriate BIN ontologies from Mercator [67], which are based on information contained in specific plant annotation databases. For doing so, we used only the genes that were differentially expressed in at least one pairwise comparison. Only DEGs with an absolute log_2_ Fold Change (LFC) greater than two were considered to facilitate the visualization of functional BIN members. Using the expression levels of these genes at 0 h as a baseline, a matrix of LFCs was obtained. Finally, we selected for visualization the diagram of biotic stress, which contains processes likely involved with the response to infection by *P. kuehnii*.

We browsed the functional annotation description fields to look for transcripts with potential relevance for plant signaling pathways or direct defense responses. Time series expression profiles were represented via line graphs and heatmaps.

### Expression analysis with qRT-PCR

We used qRT-PCR to validate the results obtained with RNA-Seq. The primers were designed with Primer3Plus [68] and Beacon Designer™Free Edition (http://www.premierbiosoft.com), with the following parameters: (i) fragment size between 90 and 200 bp; (ii) primer size ranging from 18 to 23 bp; (iii) melting temperature between 50 and 60 °C; and (iv) GC content from 40 to 60%. To verify that these primers did not spuriously amplify other transcripts, we mapped them to the transcriptome and checked that the alignments were unique.

RNA samples were treated with DNAse I (Sigma-Aldrich) and GoScript™Reverse Transcription System (Promega) was used for reverse transcription according to manufacturer instructions. qRT-PCR reactions were performed in a 7500 Fast Real-Time PCR system (Applied Biosystems) with a GoTaq® qPCR master mix kit (Promega). We used two technical replicates for each biological replicate. Reactions consisted in 6.25 *µ*L of SuperMix amplification buffer, 50 nM internal standard (ROX), 0.3 *µ*L of each primer at a 10 *µ*M, 2 *µ*L of cDNA and nuclease-free water to a final reaction volume of 12.5 *µ*L. Amplification consisted in: 95 °C for five minutes, 40 cycles of 95 °C for 10 seconds and 58 °C for 30 seconds. Product specificity was evaluated by dissociation curves. As internal controls we used a known reference gene, *glyceraldehyde-3-phosphate dehydrogenase* (GAPDH) [69], and a *Proteasome subunit beta type* that was discovered among the non-DEGs. Reaction efficiency and Ct values were determined with LinReg [70]. The relative changes in expression level were calculated with REST [71] using the same comparisons used for RNA-Seq differential expression analyses.

## Results and Discussion

### Rapid changes in gene expression in the early hours following inoculation

We sequenced libraries from 18 leaf samples collected at 0 h, 12, 24, 48 hai, 5 and 12 dai, totaling 1.17 billion reads. After quality control and trimming, 845 million reads were kept (S1 Table). The *de novo* assembly with filtered reads produced 451,462 transcripts with an N50 of 1,201 bp, and the final assembled transcriptome consisted of the 259,804 longest isoforms, with an N50 of 672 bp. On average, 67.76% of our reads were mapped to this reference. Although we expected only a minor proportion of transcript fragments from *Puccinia* in comparison with sugarcane, it is possible that their frequency increased with time as in other pathosystems [29]. Because our interest was in expression of sugarcane genes, we removed potential *P. kuehnii* transcripts. We applied homology search (BLASTX) against Swiss-Prot proteins of *Viridiplantae* and *Fungi*, allowing the identification of 219,321 probable plant transcripts, with an N50 of 671 bp, ranging from 301 bp to 15 kbp.

For differential expression analysis, we considered as expressed the genes with a CPM greater than or equal to one, in at least three samples, which resulted in 47,421 genes examined. Next, to evaluate the presence of DEGs over time, we tested contrasts between adjacent time points. Those with the largest number of DEGs were: i) 12 hai compared with 0 h: 22,148 DEGs; ii) 48 hai against 24 hai: 16,610; and iii) 12 dai compared with 5 dai: 12,438 (S1 Fig). We believe these particular time points to be of biological relevance, because spore germination in agar plates were verified at 12 hai, while microscopic observations at 24 hai confirmed fungal penetration [50]. In that case, the interval from 0 to 48 hai contains the events of spore germination, establishment of the infection and hyphae development. This shows the importance of sampling time points closer to inoculation to detect DEGs. However, it is still worthwhile to investigate the transcriptome days after inoculation, which can also reveal important information about plant metabolic responses to pathogen colonization. In this context, we note that the last sampled time point (12 dai) was marked by the confirmation of sporulation through scanning electron microscopy [50].

We found enriched terms in over and underrepresentation tests for each comparison using the BLAST2GO annotation. There were 16, 51, 48, 21 and 21 overrepresented terms for 12 hai, 24 hai, 48 hai, 5 dai and 12 dai, respectively. Similarly, we observed 12, 7, 9, 7 and 5 underrepresented terms (S1 File). In the next sections we will further explore and discuss enriched terms that help elucidate processes that were altered due to the infection. This is a fundamental step to understand temporal changes in pathways associated with disease response metabolism and plant development. Additional enrichment tests performed with the Trinotate annotation yielded similar results (S2 File), such that we will discuss them jointly, pointing occasional differences when necessary.

### Modification in pathways related to the biotic stress promoted by the *Puccinia kuehnii* infection

*P. kuehnii* infection caused significant changes in the expression of sugarcane genes involved in known molecular processes in response to biotic stresses. However, our data also suggested that the immune system was only activated two days after the inoculation (S2 File-A). We believe that *P. kuehnii* was not immediately perceived by this particular susceptible sugarcane host as a threat.

Sets of repressed DEGs were found in almost all of these processes at 12 hai (Fig 3). At this time transcripts associated with basal defenses, such as those encoding *β-1,3-glucanases*, showed changes in expression. *β-1,3-glucanases* in combination with other antifungal proteins are involved in degradation of fungal cell walls (S4 File-B and Fig 3). However, as the infection progressed more transcripts of this enzyme were detected above the basal level. The same profile was observed for transcripts involved in host cell wall reinforcements. Several genes encoding enzymes of the the *phenylpropanoid biosynthesis* pathway leading to lignin formation were identified as downregulated until 24 hai, when they were detected as upregulated (S4 FileA-D). For the GO term *response to stress* the transcripts with higher absolute LFC at 12 hai were downregulated (S2 File-C). The *immune system* was positively regulated two days after the inoculation (S2 File-A). GO term *defense response to bacterium* was among the overrepresented GO terms at 48 hai (S1 File) and *response to stress* was overrepresented at 5 dai. The latter includes, for example, transcripts encoding proteins involved in the biosynthesis of thiamine upregulated at 21 dai (S2 File-C). Thiamine is a known activator of plant disease resistance [72].

Assessment of plant hormone signaling indicated that genes related to hormones commonly involved with resistance against biotrophs, abscisic acid (ABA) and SA, were repressed at 12 hai (Fig 3). At the same time point, JA and ethylene showed roughly equal numbers of up and downregulated DEGs. Many of them later returned to the basal expression levels, similarly to what we observed for transcripts encoding transcription factors and signaling proteins.

Waves of expression, marked by downregulation followed by upregulation (or vice-versa), were evident for some groups of genes. Many coding transcripts relevant to the redox state (Fig 3 and Fig 1-D) and most of the photosynthesis/chloroplast-related genes (Fig 1-C) showed changes in opposite directions in the first (12 hai against 0 h) and third comparisons (48 against 24 hai). Initial repression of the oxidative pathways could suppress the production of reactive oxygen species (ROS). Sets of genes encoding ROS scavenging enzymes and others in *response to oxidative stress* were mostly variable in expression, with subgroups of similar expressions (S4 FileD, F-H.)

**Fig 1.**
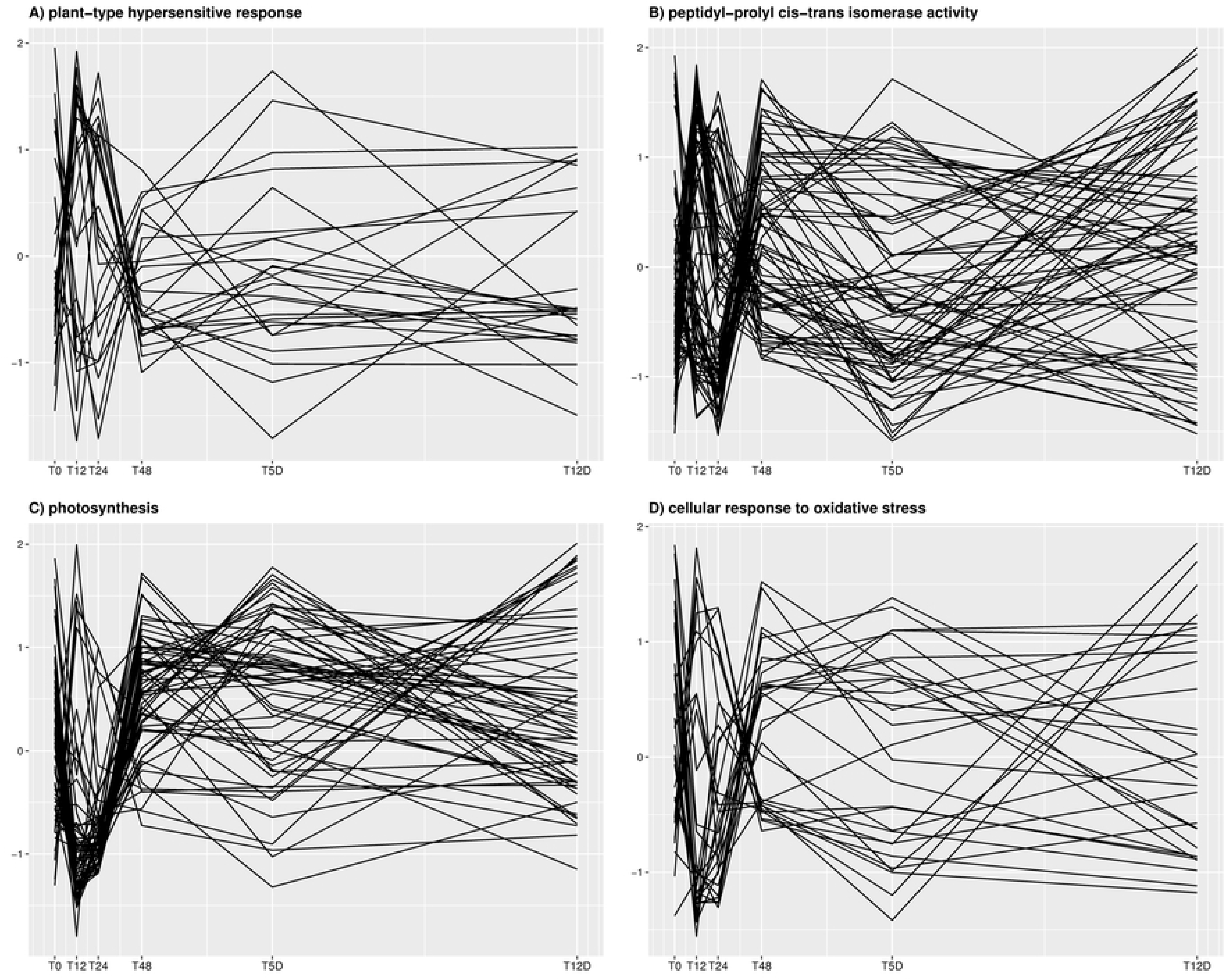
Time series profiles of differentially expressed genes (DEGs) in different Gene Ontology categories. The y-axis represents expression levels in normalized counts-per-million, and each solid black line corresponds to one DEG. Gene ontology categories are: **A)** *plant-type hypersensitive response* (GO:0009626), **B)** *peptidyl-prolyl cis-trans isomerase activity* (GO:0003755) **C)** *photosynthesis* (GO:0015979), **D)** *cellular response to oxidative stress* (GO:0034599).

### Possible late pathogen recognition by sugarcane cultivar SP89-1115

The molecular basis of plant defense mechanisms against pathogens is often explained by a two-branched immune system. The first one is based on transmembrane receptors that recognize pathogen-associated molecular patterns (PAMPs), called PAMP-triggered immunity (PTI) and the second is based on the recognition of pathogen effectors, also called effector-triggered immunity (ETI) [73–75]. Pathogen perception by the host involves the expression of resistance genes which encode proteins that typically have an NB domain and an LRR motif [74, 76], followed by the induction of the hypersensitive response, production of reactive oxygen species, cell wall reinforcement and hormone signaling [74, 75, 77].

An early fungal perception by the sugarcane plants was likely hampered because none of the GO terms usually associated with *hypersensitive response* (GO:0009626), *defense response* (GO:0006952) or *response to stress* (GO:0006950) were enriched until 24 hai (S1 File). In fact, enrichment analysis with the Trinotate annotation showed that *defense response* was significantly underrepresented within DEGs of two time points, 12 and 48 hai, and *hypersensitive response* underrepresentation started at 12 hai and persisted until 48 hai (S2 File). This indicates that in the beginning of the infection process most of these genes were not differentially expressed. It is possible that sugarcane cultivar SP89-1115 was late to recognize the molecular patterns of *P. kuehnii* to activate defense responses. Another possibility is that the plant dig recognize the PAMPs earlier, but fungal effectors may have suppressed this defense system. However, plants possibly exhibited a late response after fungal establishment, because *defense response to bacterium* (GO:0042742) transcripts were overrepresented at 48 hai (S1 File). Although transcripts of *regulation of immune response* (GO:0050776) were upregulated at 48 hours, the term was not significantly enriched. However, we found that it was overrepresented at 5 dai, because all genes were repressed at this time (S2 File and S3 File-A). Curiously, comparing 12 dai with 5 dai, the *positive regulation of defense response* was underrepresented, with no DEG annotated with this GO term (S2 File). Also, *response to stress* was enriched with DEGs at 5 dai. It is reasonable to infer that, even if late, the plant invested in defense mechanisms attempting some response.

Manual inspection of the 26 DEGs annotated with the term *defense response to bacterium* showed that in fact they represent general defense mechanisms that may act against fungi as well. They were related to protein-like kinases, heat-shock, ubiquitin and others (S3 File-B). Expression profiles of these genes did not show any clear pattern, except for a restoration to basal level at 48 hai. However, there was a small upregulated group of two *peptidyl-prolyl isomerase activity* (PPIase) enzymes and a *peroxiredoxin* at this time point. Moreover, 62.34% of the transcripts annotated with the PPIase term (GO:0003755) were differentially expressed, resulting in a significant overrepresentation (S1 File). Examining the profile of these DEGs, it is clear that they were stimulated at 48 hai (Fig 1-C). PPIases were initially associated with protein folding, but some groups are immunophilins which are involved in diverse processes [78, 79]. Cyclophilin, a protein with enzymatic PPIase activity, was accumulated in potato from 14 to 72 hours after a combination of wounding and *Fusarium solani* inoculation. The expression of this gene is influenced by the presence of ABA and JA and by abiotic stresses, in a plant tissue dependent manner [80], which agrees with the wide range of functions of the PPIases [79]. Peroxiredoxin, in turn, is a thiol peroxidase that controls redox homeostasis, photosynthesis and the transmission of redox signal [81].

The heatmap of the *defense response to bacterium* transcripts encoding a *Barwin* homolog revealed upregulation from 12 to 48 hai. The Barwin domain in plant pathogenesis-related (PR) proteins is known to be involved in a response stimulus against phytopathogenic fungi [82–84]. Another transcript, the 70 kDa *heat shock protein* (Hsp), showed significant downregulation at all time points. Members of this family are stimulated when plants are submitted to mechanical, heat or biotic stress [85]. The chaperones Hsp70 and Hsp90 trigger the expression of PR genes and defense-related transcription factors [86].

Regarding *response to stress*, most differentially expressed *calmodulin-binding* encoding transcripts (CaM-binding) were repressed at 12 hai, with an increasing expression over time (S3 File-C). Calmodulins act in transduction of Ca^+^ signaling, wherein their isoforms play a role in basal defense [87, 88]. Loss of a CaM-binding domain in an MLO (*Mildew resistance locus*) protein increased susceptibility of barley to powdery mildew fungus [89]. An upregulated CaM-binding gene was a member of the enriched *immune system* biological process in the transgenic papaya resistant to ringspot virus [90].

Five transcripts annotated with terms *thiamine* and *thiazole* were repressed at almost all times in comparison to basal levels, but notably increased at 12 dai. In fact, expression of these genes was slightly stimulated at 24 and 48 hai, but was strongly repressed at 5 dai, which resulted in the enrichment of *thiazole biosynthetic process* at this time. Thiamine has an important role in systemic acquired resistance, being known that plants treated with this vitamin can show reduced disease severity over time [72]. Infection of oil palm by *Ganoderma boninense* increased the expression of thiamine biosynthesis genes. Another study in this crop showed that four genes of thiamine biosynthesis were stimulated 24 hours after colonization by the endophytic fungus *Hendersonia toruloidea* [91]. Furthermore, both overexpression and silencing of a rice resistance gene induced a gene involved in thiamine synthesis and increased the content of this compound. Resistance genes can also balance the pathways that drive thiamine biosynthesis in chloroplasts [92]. Our results suggest that the late expression of thiamine biosynthesis genes can indicate a failure of a timely resistance signaling.

We also investigated the DEGs that composed the *hypersensitive* and *defense response* terms, where transcripts with homology to resistance genes were present. They were annotated as RGA (Resistance Genes Analogs), RPP (resistance to *Hyaloperonospora parasitica*) and RPM1 (Resistance to *Pseudomonas syringae pv. maculicola* 1) genes, which have domains commonly found in genes of the NB-LRR family [76]. Activation of resistance genes such as NB-LRR can trigger immunity by increasing PR proteins [92]. Of the 614 transcripts annotated with defense response, 102 were differentially expressed at 12 hai. At later time points the number of DEGs for this term did not exceed 70 (S3 File-D). Additionally, nearly half of the DEGs involved in *plant-type hypersensitive response* were upregulated at 12 hai, although with a low LFC (Fig 1-F). As a common result, only a small proportion of transcripts from these groups showed expression changes with high LFC and the underrepresentation is explained by the presence of many non-DEGs. In view of these results, we suggest that defense systems may not have been effective at initiating a coordinated plant response against *P. kuehnii*.

### WRKY transcription factors, cell wall reinforcement and stress-regulated transcripts altered in the first two days following inoculation

Several transcripts can be involved in avoiding pathogen establishment, encoding transcription factors, physical barrier precursors or proteins that operate directly on pathogen structure. In the first group we investigated WRKY transcription factors (S4 File-A). This is a broad class that can be found during abiotic and biotic stresses, in the latter case as regulators of defense response after infection [93]. This is achieved due to WRKY interaction with members of the mitogen-activated protein kinase (MAPK) cascade, SA and JA pathways [93–95]. WRKY-75, WRKY-90, WRKY-48 and WRKY-33 were among the most upregulated transcription factors in resistant plants of a papaya pathosystem [90]. In our study, one WRKY-33 gene was repressed at 12 hai, while two others presented expression slightly greater than zero at this moment and increased over time. Genes of WRKY-24, WRKY-45 and an uncharacterized WRKY transcription factor were upregulated at 12 hai, but more strongly at 24 and 48 hai (S4 File). An allele of WRKY-45 was reported as a negative regulator of bacterial infection in rice, while the other allele conferred resistance to plants. Interestingly, both alleles positively regulated the resistance against the fungus *Magnaporthe grisea* [96].

Disease resistance can also be initially triggered by genes that act over pathogen structure after PAMP recognition, such as *β-1,3-glucanase* and *chitinase* [73, 97]. We found two groups of *β-1,3-glucanases* with different expression patterns, particularly during the first 48 hai. The first group contained downregulated genes, with some members reaching a minimum LFC of −2.0 at 12 hai and 24 hai. The second group showed LFC values up to 5.0 at 24 hai, indicating a strong stimulus to expression (S4 File-B). These enzymes are hydrolases and are classified as PR because they act by directly degrading *β*-1,3-glucans, the most abundant polysaccharides present in fungal cell walls, and that serve as PAMPs [73, 97]. A *β-1,3-glucanase* 5 was highly upregulated at 3 hai in a peach cultivar resistant to *Xanthomonas arboricola* pv. *pruni* [34]. During smut infection in sugarcane, RNA-Seq analysis revealed an induced such gene at 48 hai. Additionally, qRT-PCR analyses revealed *β-1,3-glucanase* repression in a susceptible sugarcane cultivar, while a cultivar with external resistance exhibited upregulation [39].

A widely studied fungal PAMP is chitin, which is a structural polymer that composes part of the cell wall and is degraded by another kind of hydrolase, the *chitinase* [73]. S4 File-B shows that eight chitinase-annotated transcripts presented a pattern similar to that of the second group of *β-1,3-glucanases*, with an expression peak at 24 hai and keeping the expression above the basal level. Only three genes showed expression levels below that observed at 0 h. Others presented varying expression profiles across time points. Chitinase genes were reported as differentially expressed even in sugarcane cultivars susceptible to smut [40, 98]. A similar mixed expression pattern was also described in these cultivars, while in the case of resistance they were upregulated at 24 or 48 hai [98]. These authors also highlighted that higher expression of chitinases depends on the sampled tissue. Other studies reported chitinases as primary temporal responses in apple and peach plants infected with *Alternaria alternata* and *Xanthomonas arboricola* pv. *pruni*, respectively [38, 99], differentially expressed in coconut infected with phytoplasma [100] and in the cacao-*Moniliophthora perniciosa* pathosystem [37]. Genes involved in the synthesis of this enzyme were induced first at 1 dai and most prominently at 11 days after the inoculation of wheat susceptible to yellow rust [29].

Among the physical barriers against infection, lignin deposition in the cell wall is considered a localized response that can be hampered in susceptible plants as a result of pathogen modulation [101]. Transcripts annotated with the *lignin biosynthetic process* term (GO:0009809) showed a substantial decrease in expression in the first time point after inoculation, with some reestablishment at 48 hai (Fig 2-A). Lignin deposition in the epidermis is an important event to prevent sugarcane infection by *S. scitamineum* at 24 hai [102]. We observed that almost all genes of *cinnamyl-alcohol dehydrogenase* (CAD), a precursor of lignin biosynthesis, returned to their basal expression level after marked downregulation at 12 hai. In contrast, it is noticeable that three genes were highly upregulated from 12 hai to 12 dai (Fig 2-B). To achieve an overview of the phenylpropanoid biosynthesis, we selected DEGs annotated as enzymes of this pathway to visualize their expression profiles. Among them there were a CAD (EC:1.1.1.195) and a *caffeic acid 3-O-methyltransferase* (EC:2.1.1.68) upregulated in the transition from 12 to 24 hai (Fig 2-D). Also, all *phenylalanine/tyrosine ammonia-lyase* (EC:4.3.1.25), four *4-coumarate–CoA ligase* (EC:6.2.1.12) and all *cinnamoyl-CoA reductase* (EC:1.2.1.44) were upregulated at 48 hai. Poplar infected by a necrotroph showed temporary activation at 36 hai of genes encoding these last two enzymes [103]. Several authors noted that genes encoding enzymes of the phenylpropanoid pathway are differentially expressed in the first hours or many days after infection [33, 38, 39, 42].

**Fig 2.**
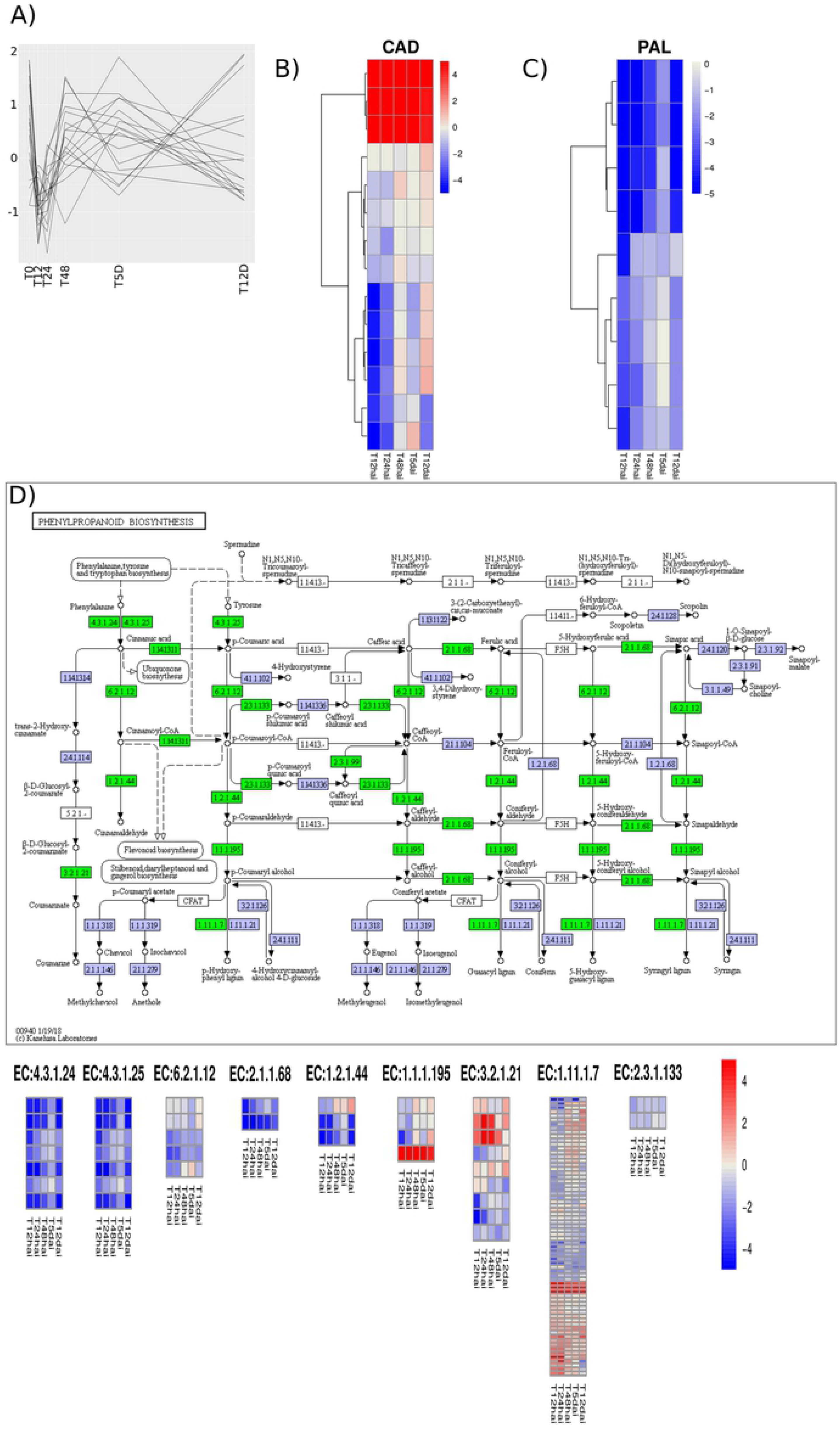
Transcripts in the phenylpropanoid biosynthesis pathway. **A)** Expression profile of transcripts annotated with term *lignin biosynthetic process* (GO:0009809); **B)** Heatmap of transcripts annotated as *cinnamyl-alcohol dehydrogenase* (CAD); **C)** Heatmap of transcripts annotated as *phenylalanine ammonia-lyase* (PAL); **D)** Phenylpropanoid biosynthesis pathway with differentially expressed Enzyme Codes (ECs) colored in green. The expression profiles of the transcripts annotated with these ECs are shown in the heatmaps at the bottom.

**Fig 3.**
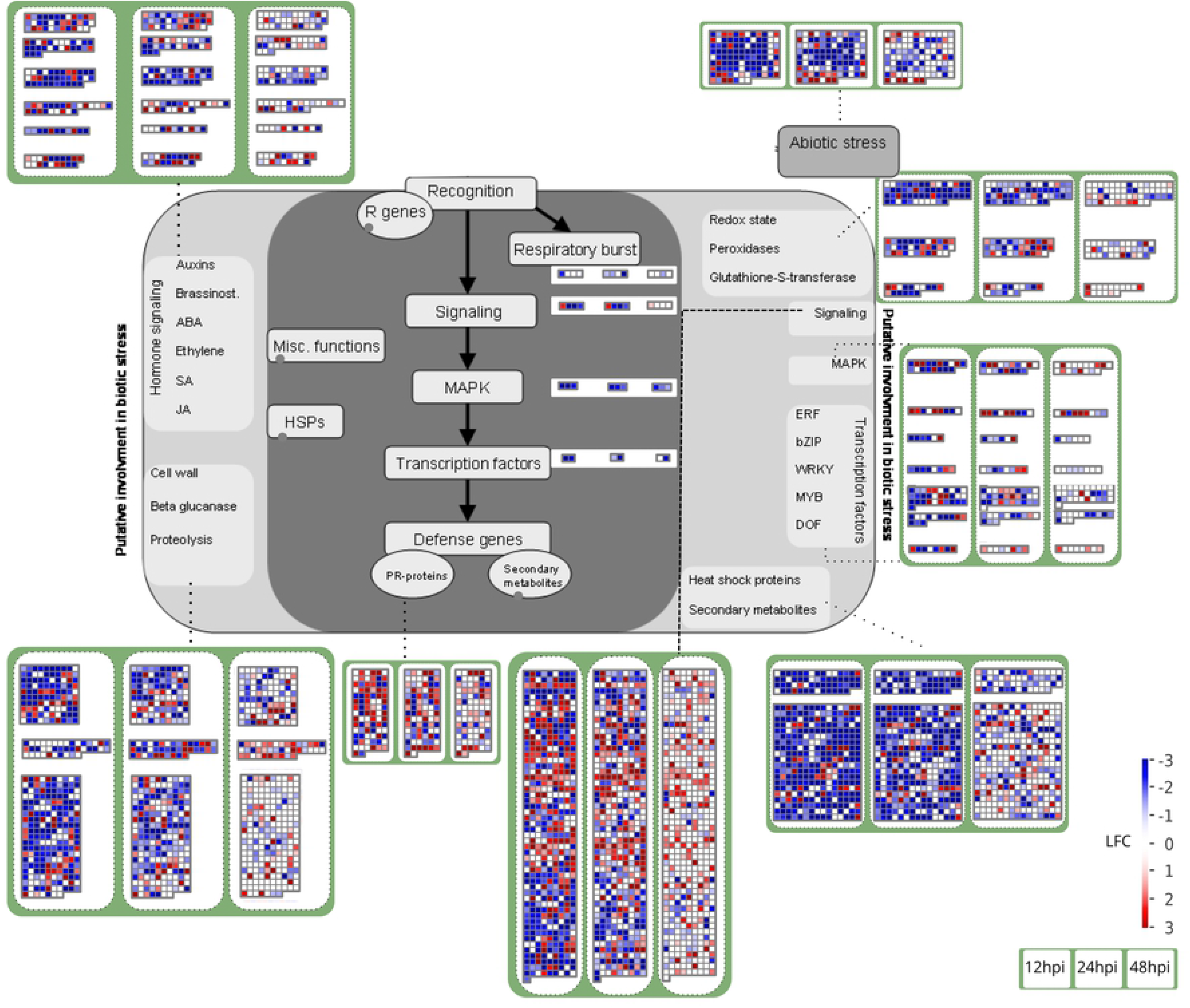
Expression profiles of transcripts mapped to processes of the biotic stress pathway. For each functional class, the three adjacent white boxes represent results for 12, 24 and 48 hai. Groups of squares in each of them represent transcripts mapped to the same process (each square is a transcript). Black dotted segments link the processes to the corresponding transcripts. The color scale indicates the log_2_ Fold Change of each transcript in comparison to the baseline expression level at 0 h.

Other authors observed that upregulation of these genes can be associated with resistance [40, 90]. In contrast, phenylpropanoid biosynthesis was enriched at 12 hai in the compatible interaction of an apple cultivar and *A. alternata* [38]. Furthermore, this pathway was enriched with downregulated genes in cucumber infected by a mutant of hop stunt viroid-grapevine at 28 dai [33]. In sugarcane smut, tyrosine accumulation is characteristically associated with phenylpropanoid biosynthesis [104]. Our results indicated that the last component of the pathway, *peroxidase* (EC:1.11.1.7), was represented by groups of genes with variable expression, as well as some with a more constant expression pattern along the investigated time interval (Fig 2-D and S4 File-D). We note that this can partly be due to the many functions of this enzyme [105].

The phenylpropanoid pathway is important in plant defense, not only by providing molecules for lignification, but also antimicrobial compounds such as coumarins and flavonoids [106]. One key enzyme that is at the start of the biosynthesis of these groups is *phenylalanine ammonia-lyase* (PAL, EC:4.3.1.25), whose accumulation has been described after elicitor stimulus and signal transduction [107]. Furthermore, this kind of response can be delayed in compatible interactions [107]. Hence, high expression of PAL-encoding genes is a common attribute of resistant cultivars [31, 34], including those of sugarcane [39, 40, 42]. In the *P. kuehnii* pathosystem the five DEGs annotated as PAL were repressed at 12 hai, but three of them were upregulated in the next hours, returning to the basal level at 5 dai (Fig 2-C). Hence we note that this delayed upregulation of PAL (Fig 2-D) was not sufficient to increase the expression levels above those observed at 0 h.

The previous processes are commonly described in the literature of plant-pathogen interactions. Investigation of other enriched terms can reveal additional transcripts potentially relevant for stress responses. We noticed that at 48 hai genes involved in transposition were significantly induced when compared with all the other time points, the same pattern as noted for *regulation of immune response*. They were annotated with the enriched term GO:0032199 (*reverse transcription involved in RNA-mediated transposition*) and were homologous to LINE-1 retrotransposons, endonucleases, exonucleases and reverse transcriptase. Enrichment with the Trinotate annotation also indicated that *transposition, RNA-mediated* (GO:0032197), viral processes (the redundant GO:0044826 and GO:0075713) and host membranes (GO:0044185 and GO:0020002) terms were enriched at 48 hai (S2 File). Activation of retrotransposons by fungal stimulus response in plants has already been described in other species [108]. Regulation of expression of transposable elements was described in sugarcane tissues [109] and expression of these elements was detected in sugarcane interacting with symbionts [110]. In rice, more than 10% of the transposable elements showed changes in expression in seedlings infected by the *Rice stripe virus*. Although most of them were downregulated at 3, 7 or 15 dai, more of these genes were upregulated as the time interval increased [30]. A long terminal repeat (LTR) retrotransposon was induced in a sugarcane genotype resistant to smut [111], and another study identified expression of a retrotransposon 24 hours after the inoculation of sugarcane mosaic virus in a resistant cultivar [112]. The infection by and development of *P. kuehnii* in sugarcane cultivar SP89-1115 did not seem to lead to a coordinated expression of these genes until 24 hai, but they were remarkably upregulated at 48 hai (S3 File-G).

### Expression waves of genes involved in photosynthesis and oxidative processes

Besides defense response and signaling transcripts, other biological processes may give indications of a possible modulation of the sugarcane metabolism by the fungus in a compatible interaction. Among these, we highlight photosynthesis and processes affecting oxidative status, which were enriched with many DEGs (S1 File). Changes in primary plant metabolism are a consequence of contact with the pathogen.

Carbohydrate metabolism is manipulated to meet the nutritional requirements of the pathogen [113]. Indeed, all *carbon utilization* (GO:0015976) genes were differentially expressed at 12 hai, along with transcripts encoding proteins of the photosynthetic apparatus. For the 89 DEGs that were annotated with photosynthesis terms (GO:0015979 and GO:0009765), 91% were downregulated at 12 hai. Later, only 32 geness were upregulated from 12 to 24 hai. Additionally, we observed that 83 genes were upregulated at 48 hai. Downregulation of genes associated with photosynthesis is a general phenomenon that can support defense response initiation against biotic stresses [113, 114], which is evidenced in RNA-Seq of infected leaves of other species [34, 37, 100]. Our data shows that this repression occurred for a majority of genes in the first 12 hours after inoculation with *P.kuehnii*, later restored to basal expression levels at 48 hai (Fig 1-C).

It is important to mention that circadian rhythm could potentially influence the observed repression of these genes at 12 hai. For that reason we also investigated the contrast between 24 hai and the basal expression level at 0 h. This comparison revealed 9,070 downregulated genes, of which 89.96% were also downregulated at 12 hai. Only 27 DEGs showed expression in inverted directions, *i.e.*, were upregulated at 12 and downregulated at 24 hai against 0 h, or vice-versa. We found 74 DEGs annotated with the photosynthesis terms at 24 hai against 0h, 90.54% of them with the same profile as at 12 hai.

Terms associated with chlorophyll, chloroplast and photosystems were overrepresented in the comparisons from 12, 24 and 48 hai (S1 File). Photosynthesis-related genes, such as the photosystem ones, can be downregulated by biotic agents [114]. Furthermore, microarray gene expression studies of *Arabidopsis thaliana* infected by *Pseudomonas syringae* revealed the repression of nuclear chloroplast genes in the early stages of infection. From 6 to 8 hai there was a decrease in photosynthetic CO_2_ accumulation, reducing net photosynthesis in comparison to mock and a non-virulent *P. syringae* strain. This occurred possibly by the action of type III effectors over photosystem II before bacterial colonization [115]. Susceptible cucumber and rice plants have also shown downregulation of chlorophyll, chloroplast and photosynthesis genes for weeks after infection [33, 116]. This is in agreement with the physiological impact of orange rust, because measures of chlorophyll content and net photosynthetic rate of sugarcane leaves decrease according to susceptibility to *P. kuehnii* [18].

The *oxidation-reduction* (GO:0055114) term was enriched at 12, 24 and 48 hai and, later, at 12 dai. This process together with *oxidoreductase activity* (GO:0016491 and GO:0016705) had 596 downregulated genes at 12 hai and 468 upregulated at 48 hai. Evidence of an oxidative imbalance is complemented by enrichment of *redox homeostasis* (GO:0045454) at 12 and 24 hai, with redoxins mostly downregulated in the former and upregulated in the latter time. Reactive oxygen species (ROS) produced by these metabolic pathways act as propagating signal during stresses, with the advantage of their fast production and scavenging. They can trigger defense against phytopathogenic fungi due to their role in signal transduction, localized cell death and modifications to molecules, like proteins and lipids, and to the cell wall [117–120]. Infection by *Acidovorax avenae* subsp. *avenae* resulted in enrichment of oxidation-reduction pathways with up and downregulated sugarcane genes [42].

We note that oxidative pathways are possibly connected with some of the previously presented results. For example, presence of ROS is one potential reason for Hsp70 accumulation [85] and for the activity of calmodulins, amine oxidases and other types of peroxidases [117]. A peroxiredoxin of defense response to bacterium was stimulated at 48 hai, possibly as a result of the presence of ROS, because it is a sensor peroxidase [81]. Amine oxidases collaborate in ROS production and are upregulated in resistant apple and peach plants submitted to biotic stress [34, 121]. These genes formed a large group characterized by repression at 12 hai, with some members showing an LFC less than −2.0 until 24 hai (S4 File-E). ROS generation in the periphery of cells is a complex phenomenon that can represent a plant response by oxidative cross-link of cell walls and defense signaling [20]. Oxidative burst usually follows gene-for-gene resistance and ROS production is important for hypersensitive cell death [23]. Given that the *plant-type hypersensitive response* was underrepresented, we can suppose that some of the oxidation-reduction proteins could not act to trigger hypersensitivity. Suppressing the oxidative burst may be strategical to the fungus because ROS have a localized action promoting cell death, and a biotroph needs live tissues to colonize [20, 23]. In summary, products of transcripts annotated with oxidative terms in this pathosystem possibly failed to activate additional pathways that establish host defense.

Other pathways are also associated with the production of ROS. For instance, a decrease in photosynthesis can be associated with repression of defense signaling. This happens due to effector inhibition of photosynthetic electron transport, which restricts ROS production leading to suppression of PTI [115]. Some ROS scavenging enzymes that are targeted to the chloroplast are likely downregulated for ROS direct or indirect defense [114]. ROS signaling hypotheses suggest that stimuli create waves requiring other signals or that specific ROS waves are perceived by other mechanisms, or even that ROS waves are transferred from stress receptors to response targets [120].

Pathways of oxidative detoxification are also relevant in defense response signaling [31]. The expression profile of genes involved in the cellular response to oxidative stress was similar to that of photosynthesis (Fig 1-D), with roughly 70% of their genes being differentially expressed at 48 hai (S1 File). The term *response to oxidative stress* (GO:0006979) was enriched at the same time point, but detailed investigation showed a similar number of up and downregulated DEGs. This term was composed to a large extent by peroxidases (S3 File-E). Redox homeostasis is also achieved by other enzymes involved in ROS scavenging, such as catalases (CAT), superoxide dismutases (SOD) and glutathione transferases (GST) [117]. In this work, we did not observe a clear expression pattern over time for these genes, because they formed small groups with differing profiles (S4 File-F-H).

### Validation of differential expression with qRT-PCR

We selected transcripts for validation based on the differential expression analysis results and on their biological relevance to the infection process. Some of them showed significant differential expression in all tested comparisons: *probable cinnamyl alcohol dehydrogenase 8D*, *calmodulin binding protein2*, *putative ethylene response sensor 2* and *glutathione transferase*. Others were differentially expressed on two to three comparisons: *catalase*, *probable WRKY transcription factor 70*, *Superoxide dismutase*, *small heat shock protein*, *Glucan endo-1,3-beta-glucosidase*, *bZIP transcription factor* and *chloroplast ribulose-1,5-bisphosphate carboxylase/oxygenase small subunit*. The only non-DEG selected was a *Proteasome subunit beta type*, used as internal control along with GAPDH. Each primer pair was designed to ensure effective amplification of a unique transcript and sequences are available in the S2 Table. Overall, we performed 55 tests: five adjacent time comparisons for eleven transcripts. Of these, only seven did not confirm the RNA-Seq differential expression results. This demonstrated good correspondence between RNA-Seq and qRT-PCR results (S2 Fig).

Even though the number of matching results was high between the two techniques, it is important to point out some details. There were two cases when they were discordant: i) Detection of significance with only one technique; ii) Significant changes in opposite directions. For the first case, RNA-Seq seems to have shown more power to detect significance for small Fold Changes, as seen for BZIP at 12 hai, SOD and GST at 24 hai. On the other hand, we detected significance for GLU at 48 hai only with RT-qPCR. This can be explained because the CPMs of this transcript were highly variable between biological replicates, resulting in higher residual variance. The second case was verified for HEAT at 48 hai, with RNA-Seq indicating significant downregulation and RT-qPCR showing upregulation of this transcript. Both the forward and reverse primers aligned to four isoforms of this transcript, but only the forward primer aligned against the longest isoform. There is a possibility that these primers amplified other *heat shock* transcripts. Because RNA-Seq and RT-qPCR are methods with different properties, it was expected that not all LFCs were in agreement for both.

### Successful infection of susceptible sugarcane

A biotrophic fungus typically germinates on the leaf surface, forms appressoria to invade the host through the stomata, feeds from the host cells by its haustoria and reproduces itself afterwards by producing numerous uredospores [122]. The infection can be avoided by recognition of pathogen patterns in PTI inducing cellular responses. A second mechanism, ETI, occurs when proteins of resistance genes operate on virulence proteins of the pathogen [74]. A viable strategy to capture the dynamics of these molecular events is to detect changes in gene expression according to the progress of plant infection [31]. In the sugarcane-*P. kuehnii* pathosystem, appressoria and pustules were observed by Gomez *et al.* [50] at 24 hai and 12 dai, respectively, and symptoms were detected at 9 dai, followed by sporulation two days later. Our analyses revealed certain genes upregulated in the beginning of the infection process. Among these, we highlight genes of the lignin pathway, some chitinases and PR genes. Other genes were repressed, such as representatives of CaM-binding and photosynthesis-related proteins. Importantly, we observed underrepresentation of members of hypersensitive and defense responses in functional enrichment analysis. A delayed defense response by cultivar SP89-1115 was suggested by the upregulation of some genes present in groups of PPIases, *β*-1,3-glucanase, thiamine-thiazole biosynthesis, PALs and CADs.

The global impact of the biotroph *P. kuehnii* on sugarcane was that for most processes of the biotic stress pathway, genes downregulated at 12 hai gradually had their expression increased back to the basal level (Fig 3). However, we noted that some PR-protein annotated transcripts were upregulated, opening up the possibility of some early plant responses. This is also reinforced by the upregulation of a part of the larger group of signaling transcripts. We do not rule out the existence of genes regulated to favor the fungus. As for hormone metabolism, it is possible to note that the majority of genes associated with SA were initially downregulated and then returned to the initial expression at 48 hai. This result is expected because this plant hormone is decisive to establish defense signaling against many biotrophs [20, 23]. Biotrophic pathogens may act by modulating the signaling pathways to avoid cell death, keeping the host cell alive as long as necessary [123]. The biotrophic pathogenic fungus *Ustilago maydis* secretes the chorismate mutase (Cmu) into the plant cell, modulating the SA pathway during the infection in maize [124]. Fungal Cmu1 interferes in the chorismate metabolism to produce phenylalanine or tyrosine via the prefenate precursor instead of isochorismate [125]. Genes related to the metabolism of another hormone, ABA, were downregulated at 12 hai and remained repressed at 24 hai (Fig 3). Indeed, enrichment of *abscisic acid homeostasis* was verified in the contrast between 24 and 12 hai (S2 File). The role of ABA in plant-pathogen interaction is that its biosynthesis and signal transduction are potentially involved with partial resistance, as detected in *Avena sativa* infected by *Puccinia coronata* [31]. ABA signaling was verified by ten ABA genes being upregulated in papaya infected by ringspot virus, while only two were downregulated [90].

A study in two poplar cultivars infected with the hemibiotrophic fungus *Marssonina brunnea* found *response to external stress* enrichment in both susceptible genotypes only at 96 hai, the last investigated time period after infection. Stress-related genes quickly showed changes in expression, manifested as a general downregulation pattern of an Hsp at 6 hai [103]. McNeil and coworkers [39] detected sugarcane response against *S. scitamineum* at 48 hai in an incompatible interaction. However, they did not exclude the possibility of it occurring earlier, in a not sampled time point. Differentially expressed genes in sugarcane infected by this pathogen were found in greater number at earlier sampled time points in resistant cultivars [40]. Early responses may be necessary to initiate a successful defense strategy, which seems not to have been the case in the particular interaction we studied. In addition, we noted changes in transcriptional profiles that can reveal modification of plant metabolism to favor the fungus. In this case, it is possible that not only the plant was unable to respond quickly, but that *P. kuehnii* effectively disabled the existing defense mechanisms after initial contact.

Finally, investigation of time series data creates the possibility of exploring annotated transcripts and pathways from both host and pathogen, observing significant changes in gene expression. From the host perspective it is also possible to find differences between resistant and susceptible cultivars [29], or as a consequence of varying responses against particular pathogen strains [103]. Differences between genotypes are a consequence of genes being expressed at different levels in some genotypic backgrounds, and also due to the timing of expression of resistance genes, which can be delayed in susceptible plants [31]. Our research of sugarcane infected by *P. kuehnii* pointed that the earlier time points likely have great importance to understand possible defense responses. Also, it yields new hypotheses for future research with the aim to improve our knowledge of possible sugarcane defense pathways to avoid orange rust.

## Conclusion

Profiles of differentially expressed genes over time series data, grouped by similar metabolic functions, provided an overview of how sugarcane transcripts were influenced by inoculation with *P. kuehnii*. Most DEGs were found at the beginning of the infection process, at 12 hai, as well as during fungal growth in leaf tissue and after apparent symptoms and sporulation. We detected as differentially expressed a portion of the transcripts functionally related to disease response, and some groups presented the same expression profiles in specific time comparisons. Other enriched processes contained transcripts with a coordinated expression pattern, such as the photosynthesis-related ones, or revealed several groups with diverse patterns, as seen in the oxidation-reduction pathways. This analysis also offered the opportunity to examine some of the transcripts involved in pathogen recognition and immune response processes. Together, these observations revealed increased expression over time of genes related to PAMP recognition and signaling. The pathogenic agent may have acted directly on host plant metabolism by modulating particular pathways. Yet, this does not exclude the possibility of a delayed sugarcane response, mainly from 24 to 48 hai. In this context, additional studies can explore the expression profiles of the detected transcripts in sugarcane cultivars with distinct degrees of resistance to *P. kuehnii*.

### Database Accession Number

The sequencing data has been deposited at DDBJ/EMBL/GenBank under the BioProject ID PRJEB31605.

## Supporting information

**S1 Table. Sequencing libraries of sugarcane cultivar SP89-1115 infected with *Puccinia kuehnii*.** The columns contain sample identification (the sampled time point and replicate), the number of raw reads, the percentage of processed reads and the overall alignment rate.

**S1 Fig. Distribution of non-differentially expressed, down and upregulated genes for each time comparison.** Each bar indicates a different time contrast between adjacent time points. Up and downregulated DEGs are indicated by the colors red and blue, respectively. Non differentially expressed genes (Non-DEG) are indicated by the gray color. The time points of 5 dai and 12 dai were written as 5d and 12d.

**S1 File. Tables of significantly overrepresented and underrepresented gene ontology (GO) terms in each time comparison, based on the BLAST2GO functional annotation.** For each significant GO term, the columns indicate: Category, Description, corresponding *p*-value of the test, the total number of transcripts annotated in the category and the number of DEGs. Every two tables represent the over and underrepresented GO terms, respectively, for each time point comparison, according to the following order: (*A-B*) 12 hai compared to 0 h, (*C-D*) 24 compared to 12 hai, (*E-F*) 48 compared to 24 hai, (*G-H*) 5 dai compared to 48 hai, (*I-J*) 12 compared to 5 dai.

**S2 File. Tables of significantly overrepresented and underrepresented Gene Ontology (GO) terms in each time comparison, based on the Trinotate functional annotation.** For each significant GO term, the columns indicate: Category, Description, corresponding *p*-value of the test, the total number of transcripts annotated in the category and the number of DEGs. Every two tables represent the over and underrepresented GO terms, respectively, for each time point comparison, according to the following order: (*A-B*) 12 hai compared to 0 h, (*C-D*) 24 compared to 12 hai, (*E-F*) 48 compared to 24 hai, (*G-H*) 5 dai compared to 48 hai, (*I-J*) 12 compared to 5 dai.

**S3 File Heatmaps of differentially expressed genes (DEGs) in Gene Ontology (GO) enriched terms.** Each row represents a DEG annotated with the corresponding GO Term. Columns indicate the time points after inoculation, ranging from 12 hai to 12 dai. Cell values represent the log_2_ Fold Changes between the five times points in comparison to the baseline expression level at 0 h, truncated at −5.0 and 5.0. **A)** *Defense response to bacterium*, **B)** *Regulation of immune response*,**C)** *Response to stress*, **D)** *Defense response*, **E)** *Response to oxidative stress*, **F)** *Plant-type hypersensitive response*, **G)** Transposition-associated terms.

**S4 File. Heatmaps of differentially expressed genes (DEGs) with manually investigated annotations.** Each row represents a DEG and columns indicate the time points after inoculation, ranging from 12 hai to 12 dai. Cell values represent the log_2_ Fold Changes between the five times points in comparison to the baseline expression level at 0 h, truncated at −5.0 and 5.0. Terms were searched with the following descriptions: **A)** *WRKY*, **B)** *β-1,3-glucanase*, **C)** *chitinase*, **D)** *peroxidase*, **E)** *amine oxidase*, **F)** *catalase*, **G)** *superoxide dismutase*, **H)** *glutathione transferase*.

**S2 Table. Sequences of primers used for qRT-PCR validation based on differentially expressed genes.**

**S2 Fig. Expression comparison between RNA-Seq and RT-qPCR.** The *y*-axis contains the log_2_ Fold Change for the comparisons in the *x*-axis. RNA-Seq (RT-qPCR) is shown in green (blue). Significant contrasts are indicated with an asterisk, while non-significant ones are marked as ns. Each graph corresponds to a different transcript: **A)** *putative ethylene response sensor 2*, **B)** *catalase-1*, **C)** *probable WRKY transcription factor 70*, **D)** *Superoxide dismutase*, **E)** *small heat shock protein*, **F)** *glucan endo-1,3-beta-glucosidase 12*, **G)** *calmodulin binding protein2*, **H)** *bZIP transcription factor*, **I)** *chloroplast ribulose-1,5-bisphosphate carboxylase/oxygenase small subunit*, **J)** *probable cinnamyl alcohol dehydrogenase 8D*, **K)** *glutathione transferase*.

## Acknowledgments

We thank the sequencing facility Center of Functional Genomics (USP, Campus “Luiz de Queiroz”); Laboratório de Genética Molecular, Departamento de Fitopatologia e Nematologia (USP, Campus “Luiz de Queiroz”) and Laboratório de Biotecnologia de Plantas (UFSCar, Centro de Cîencias Agŕarias).

